# Gene-specific RNA homeostasis revealed by perturbation of coactivator complexes

**DOI:** 10.1101/2024.01.30.577960

**Authors:** Faezeh Forouzanfar, Damien Plassard, Audrey Furst, David F. Moreno, Karen A. Oliveira, Bernardo Reina-San-Martin, László Tora, Nacho Molina, Manuel Mendoza

**Affiliations:** Institut de Génétique et de Biologie Moléculaire et Cellulaire, Illkirch, France; Centre National de la Recherche Scientifique, UMR7104, Illkirch, France; Institut National de la Santé et de la Recherche Médicale, U964, Illkirch, France; Université de Strasbourg, Strasbourg, France

## Abstract

Transcript buffering entails the reciprocal modulation of mRNA synthesis and degradation rates to maintain stable RNA levels under varying cellular conditions. Current research supports a global, non-sequence-specific connection between mRNA synthesis and degradation, but the underlying mechanisms are still unclear. In this study, we investigated changes in RNA metabolism following acute depletion of TIP60/KAT5, the acetyltransferase subunit of the NuA4 transcriptional coactivator complex, in mouse embryonic stem cells. By combining RNA sequencing of nuclear, cytoplasmic, and newly synthesised transcript fractions with biophysical modelling, we demonstrate that TIP60 predominantly enhances transcription of numerous genes, while a smaller set of genes undergoes TIP60-dependent transcriptional repression. Surprisingly, transcription changes caused by TIP60 depletion were offset by corresponding changes in RNA nuclear export and cytoplasmic stability, indicating gene-specific buffering mechanisms. Similarly, disruption of the unrelated ATAC coactivator complex also resulted in gene-specific transcript buffering. These findings reveal that transcript buffering functions at a gene-specific level and suggest that cells dynamically adjust RNA splicing, export, and degradation in response to individual RNA synthesis alterations, thereby sustaining cellular homeostasis.

## Introduction

Eukaryotic gene expression involves a precise sequence of events, starting with the synthesis of messenger RNA (mRNA) precursors in the nucleus. Subsequently, these molecules are processed, spliced and exported to the cytoplasm, where they undergo translation into proteins before ultimately undergoing degradation. Initially studied as distinct RNA metabolic processes, it is increasingly evident that RNA synthesis and degradation are inherently linked. In particular, the development of new methods to measure mRNA transcription and degradation rates uncovered connections between nuclear mRNA synthesis and cytoplasmic degradation. For instance, global reduction of transcription rates leads to a corresponding increase in mRNA stability in budding yeast (Baptista et al., 2017; Sun et al., 2012; Warfield et al., 2017) and in animal cells (Berry et al., 2022; Helenius et al., 2011). Conversely, global inhibition of mRNA degradation is associated with decreased transcription rates (Sun et al., 2012, 2013; Haimovich et al., 2013). This phenomenon, termed “transcript buffering”, is thought to maintain mRNA concentration constant, which may be important in physiological contexts such as during changes in cell size (Timmers and Tora, 2018; Hartenian and Glaunsinger, 2019; Berry and Pelkmans, 2022).

Although the molecular mechanisms responsible for transcript buffering are unclear, proposed models include negative feedback on RNA polymerase II activity exerted by mRNA degradation factors (in yeast) or nuclear RNA (in animal cells) (Sun et al., 2013; Berry et al., 2022). Importantly, these models aim to explain global buffering, wherein overall changes in mRNA synthesis are compensated by overall changes in mRNA stability, and vice-versa. Yet, whether and how transcription buffering also occurs in a transcript-specific manner remains unclear.

The Tip60 complex, also known as NuA4, is an evolutionarily conserved lysine acetyl-transferase (KAT) and chromatin remodeler complex with roles in transcription, DNA damage response and intracellular signalling (Doyon and Côté, 2004; Squatrito et al., 2006). Its KAT subunit TIP60, or KAT5, is essential for early mouse development. Knockdown of *Tip60* in mouse embryonic stem cells (ESCs) leads to their reduced proliferation, loss of pluripotency, and alteration of mRNA levels (Fazzio et al., 2008).

Given the strong correlation between transcription and acetylation of histones and chromatin-associated proteins (Shvedunova and Akhtar, 2022), TIP60 would be expected to stimulate transcription. Surprisingly, RNA sequencing (RNA-seq) experiments in mESCs revealed that although the level of some mRNAs is reduced after downregulation of *Tip60*, the majority of affected mRNAs actually become more abundant. This led to the proposal that TIP60 might function predominantly as a transcriptional repressor in ESCs (Fazzio et al., 2008; Chen et al., 2013). An alternative explanation, however, is that the observed increase of mRNA levels in TIP60-deficient cells could reflect post-transcriptional dysregulation. Indeed, RNA-seq measures RNA steady-state abundance, which depends on both RNA synthesis and degradation rates. Notably, the budding yeast homologue of TIP60 has been shown to promote both the synthesis and nuclear export of mRNA (Gomar-Alba et al., 2022), suggesting that TIP60 may also regulate gene expression post-transcriptionally. Therefore, characterisation of multiple steps in RNA metabolism is essential to understand the role of TIP60 in gene expression regulation.

Here, we investigated the effects of acute depletion of TIP60 on mRNA metabolism in mouse ESCs by deep sequencing of newly synthesised, nuclear and cytoplasmic RNA. This showed that TIP60 acts mainly as a transcriptional activator, crucial for mRNA synthesis of its target genes. Only a small proportion of genes increased their transcription after TIP60 depletion, which might have occurred via indirect mechanisms. Strikingly, the observed changes in the transcription rates following TIP60 depletion were mirrored by opposite changes in RNA export and stability in both the nucleus and cytoplasm. Notably, the degree of this buffering effect corresponded precisely to the magnitude of transcriptional changes. These findings indicate that transcript buffering operates at the gene-specific level rather than globally, with important implications for our understanding of transcript buffering mechanisms.

## Results

### Loss of TIP60 halts proliferation of mouse embryonic stem cells

We established a conditional degradation system designed for the rapid depletion of TIP60 upon auxin addition in mouse ESCs. Utilising CRISPR/Cas9 gene editing, we knocked in sequences encoding an auxin-inducible degron (AID) and BioTag into both *Tip60* alleles within a ESC line expressing Tir1 (Fischer et al., 2021), which drives auxin-dependent ubiquitination of AID proteins. The resulting cell line (*Tip60^AID^*) was validated by PCR (Supplementary Figure **S1**). Western blot analyses showed that the TIP60-AID protein was efficiently eliminated within the first 6 hours of auxin treatment, and its loss was sustained for >2 days (Figure **1A**).

**Figure 1:**
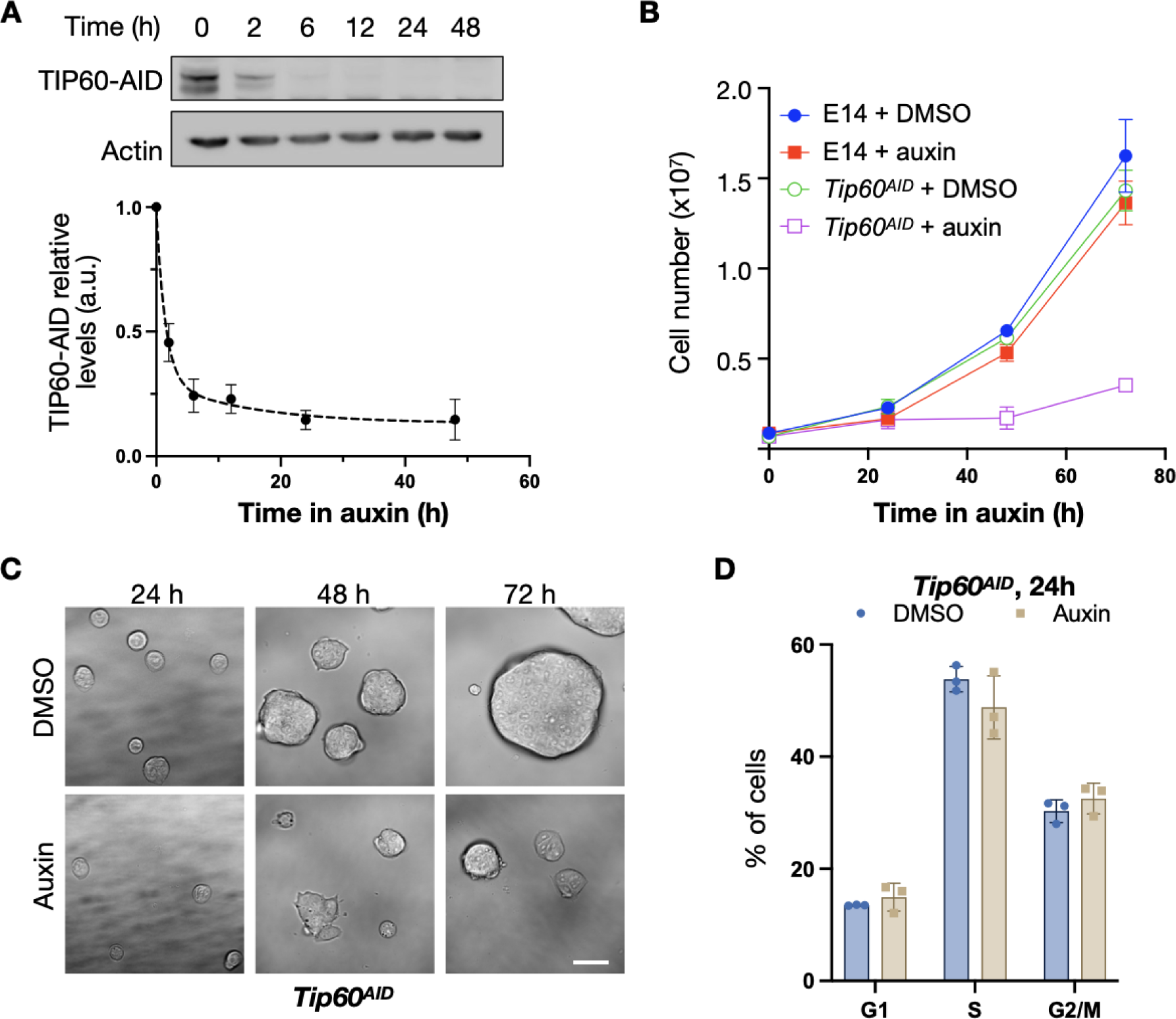
TIP60 is essential for mESCs proliferation. **A**: *Tip60^AID^* cells were incubated for the indicated time with either DMSO or 1 mM auxin. TIP60 was detected by western blotting using HRP-coupled streptavidin. Actin was used as a loading control. A representative western blot and quantification of three independent replicates (mean and standard error, SEM) are shown. **B**: Cell number (mean and SEM of n=3 independent experiments) of the indicated cells treated with DMSO or 1 mM auxin at the indicated times. **C:** Brightfield images of cells incubated with DMSO or 1 mM auxin for the indicated times. Scale bar, 40 μm. **D**: Cell cycle distribution (mean and standard deviation of three independent experiments) of *Tip60^AID^* cells treated with DMSO or auxin for 24 h, determined by flow cytometry. Cells were grown in FCS + LIF + 2i.

To evaluate the impact of TIP60 depletion on cell survival, we examined *Tip60^AID^* cell proliferation under continuous auxin exposure for 24, 48 and 72 hours. These cells were cultured in the presence of leukaemia inhibitory factor (LIF), and two inhibitors targeting mitogen-activated protein kinase (MEK)/extracellular signal-regulated kinase (ERK) and glycogen synthase kinase 3 beta (GSK3b) pathways (LIF+2i medium), which sustains uniform expression of pluripotency network genes (Hastreiter et al., 2018). In this medium, *Tip60^AID^* cells exhibited proliferation comparable to cells expressing untagged TIP60 (E14). However, the addition of auxin specifically halted proliferation of *Tip60^AID^* cells after 24 hours (Figure **1B-C**). Additionally, we observed that *Tip60^AID^* cells cultured in the presence of LIF but without 2i (permitting heterogeneous expression of pluripotency genes) proliferated at a slower rate than ESCs expressing untagged TIP60. Remarkably, auxin halted growth of *Tip60^AID^* cells even in these conditions (Supplementary Figure **S2**). These results indicate that TIP60 is essential for stem cell proliferation.

### Tip60 promotes transcription of its target genes

To directly determine the role of Tip60 in RNA synthesis, we used Transient Transcriptome sequencing (TT-seq) (Schwalb et al., 2016). TT-seq, which measures newly synthesised RNA, involved a 10-minute incubation with 4-thiouridine (4sU) to label nascent RNA in *Tip60^AID^* ESCs treated with DMSO or auxin to deplete TIP60. Subsequently, nascent RNA was fragmented, purified and quantified by Illumina sequencing. To ensure global normalisation of TT-seq data, we spiked in labelled RNA from *Drosophila*. TT-seq samples were enriched in intronic reads relative to RNA-seq samples, consistent with reproducible and efficient isolation of newly synthesised transcripts (Supplementary Figure **S3**). We conducted three independent sets of RNA-seq and TT-seq experiments utilising *Tip60^AID^* cells cultured in LIF+2i medium, treated with auxin or DMSO for 24 hours. This duration was selected to ensure a complete Tip60 depletion in auxin-treated cells without detectable perturbation of cell growth or cell cycle progression (**Figure 1, A-D**).

We compared data obtained with conventional RNA-seq with TT-seq for *Tip60^AID^* cells treated separately with auxin and DMSO. We classified RNAs exhibiting a greater than two-fold change between the DMSO and auxin conditions, with a significance level of Benjamini -Hochberg-adjusted p <0.05, as differentially expressed genes (DEGs). In RNA-seq experiments, a total of 1542 DEGs were identified, with 70% displaying upregulation (1075 mRNAs) and only 30% were exhibiting downregulation (467 mRNAs) (Figure **2A**). Downregulated genes exhibited significant enrichment for TIP60 target genes, as identified by chromatin immunoprecipitation of TIP60 (Ravens et al., 2015). Conversely, the upregulated genes were less likely to interact with TIP60 directly (Figure **2B**). Upregulated genes included factors associated with multicellular development Gene Ontology (GO) terms (Figure **2C**). These results are in line with previous *Tip60* siRNA knockdown experiments which suggested that TIP60 represses differentiation genes (Fazzio et al., 2008; Acharya et al., 2017).

**Figure 2:**
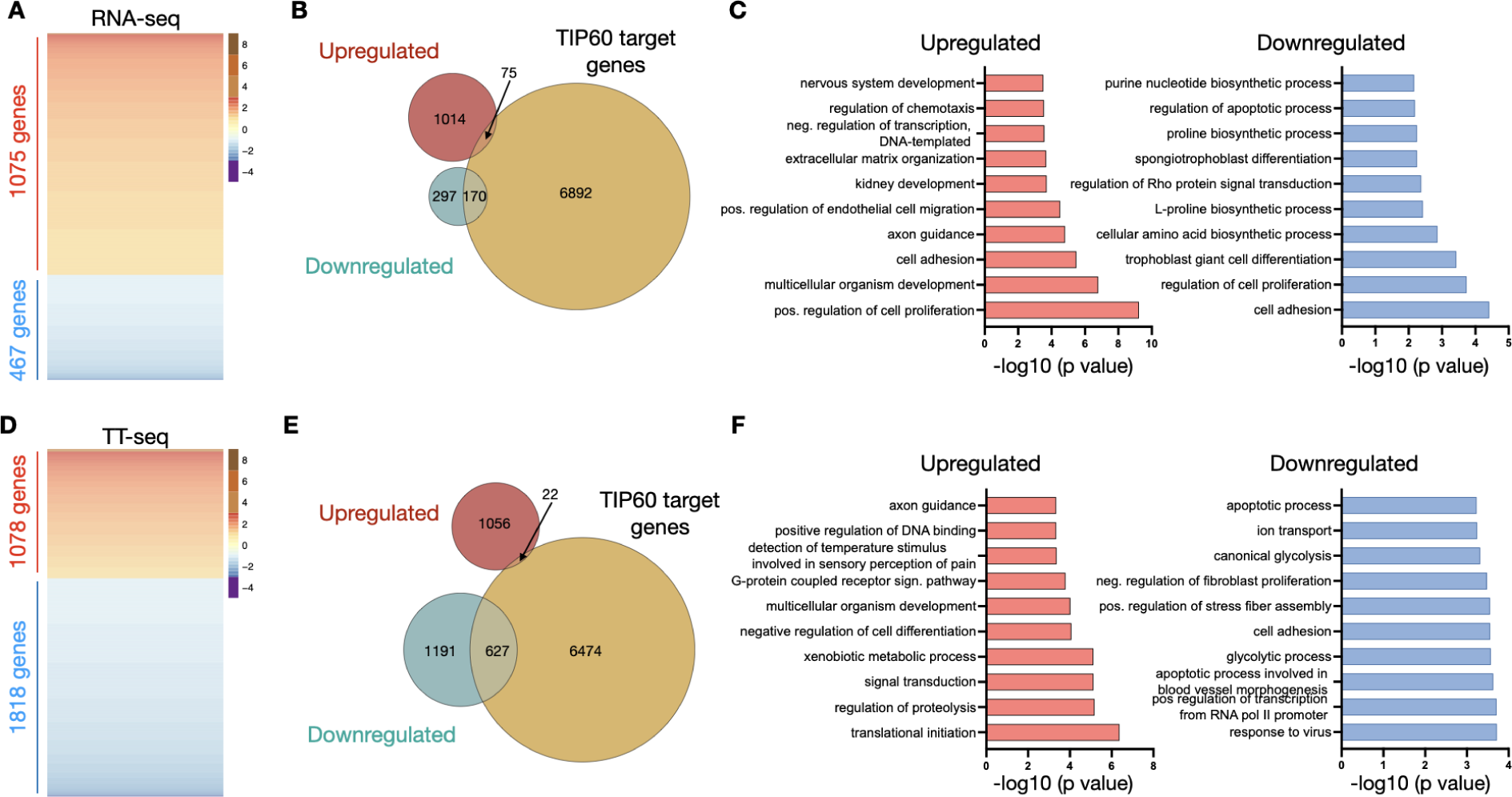
Role of TIP60 in mRNA synthesis. **A**: Heat maps of differentially expressed genes in *Tip60^AID^* cells treated with auxin vs DMSO for 24 h assessed by RNA-seq. Genes in the heatmaps are sorted from the most upregulated to the most downregulated genes. **B**: Venn diagrams designating the overlap between differentially expressed genes in (A) and TIP60-associated genes (Ravens et al., 2015). **C**: Top 10 gene ontology terms (Biological Process) associated with differentially expressed genes in (A), ranked by p-value. **D-F**: Heat maps (**D**), Venn diagrams (**E**) and gene ontology analysis (**F**) of differentially transcribed genes assessed by TT-seq. Genes were considered significantly misregulated if their log2 (fold change) was > 1 or <-1, and their Benjamini-Hochberg adjusted p value < 0.05 (n=3 independent experiments).

Intriguingly, analysis of the TT-seq data showed that TIP60 depletion had a clearly distinct effect on newly synthesised transcripts, with >60% of DEGs showing a decrease (1818 out of 2896 RNAs) and <40%, an increase in transcription efficiency (1078 RNAs) (Figure **2D**). Transcriptionally downregulated genes were specifically enriched for genes associated with TIP60 (Figure **2E**) and the TIP60 complex component p400 (Chen et al., 2015) (Supplementary Figure **S4**), while those exhibiting increased transcription included TIP60-independent developmentally regulated genes (Figure **2F**).

In summary, TIP60 depletion predominantly results in reduced transcription of its target genes. This indicates that TIP60 likely functions as a transcriptional co-activator for these genes. Importantly, this role is masked in total RNA-seq data, possibly due to compensatory changes in mRNA stability.

### Integration of TT-seq and Frac-seq data reveals gene-specific buffering

The above results suggest that the depletion of TIP60 affects more than one step of the gene expression process. To estimate how TIP60 depletion impacts the rates at which RNAs flow from the nucleus to the cytoplasm and the relative stability of RNAs in these cellular compartments, we combined TT-seq with Fractionation sequencing (Frac-seq) (Lee et al., 2020a). Frac-seq allowed to determine the fold change of RNA isolated from the nuclear and cytoplasmic compartments in control versus TIP60-depleted cells, under the same conditions as the TT-seq experiments. Validation via reverse transcription and quantitative PCR (RT-qPCR) confirmed the absence of cross-contamination between nuclear and cytoplasmic fractions. The nuclear fractions were enriched in introns and nuclear long noncoding RNAs (lncRNAs), whereas mature mRNAs were predominantly found in the cytoplasmic fractions (Supplementary Figure **S5**).

First, we used Frac-seq data to calculate the fold change of RNA isolated from the nucleus and cytoplasm in control versus TIP60-depleted cells. Overall, TIP60 depletion did not lead to a global accumulation of RNAs in either compartment. However, we detected slight changes in nucleus / cytoplasmic (N/C) ratio for specific RNA types, such as mRNAs encoding intronless genes (primarily canonical histones) and ribosomal proteins (Supplementary Figure **S6**), and long (>50 kb) lncRNAs (Supplementary Figure **S7**). Nonetheless, these changes were relatively small (less than two-fold). Thus, TIP60 depletion has a modest impact on the N/C distribution of RNAs.

Next, we integrated the TT-seq and Frac-seq datasets to fit a biophysical model that explicitly describes the following processes involved in RNA metabolism: transcription, splicing, nuclear export, and cytoplasmic degradation. This model extends the RNA velocity approach employed for analysing spliced and unspliced RNA reads from bulk and single-cell RNA-seq experiments (Gaidatzis et al., 2015; La Manno et al., 2018) to include the translocation between nuclear and cytoplasmic compartments. Furthermore, we assumed that the 24-hour auxin treatment duration was sufficiently long relative to the typical mRNA half-life, which is on the order of hours (Schwanhäusser et al., 2011; Sharova et al., 2009), allowing cells to attain a new quasi-equilibrium state after the perturbation. Consequently, we solved the model for the steady state, simplifying the fitting procedure (see Figure **3A** and *Methods* for extended details).

**Figure 3:**
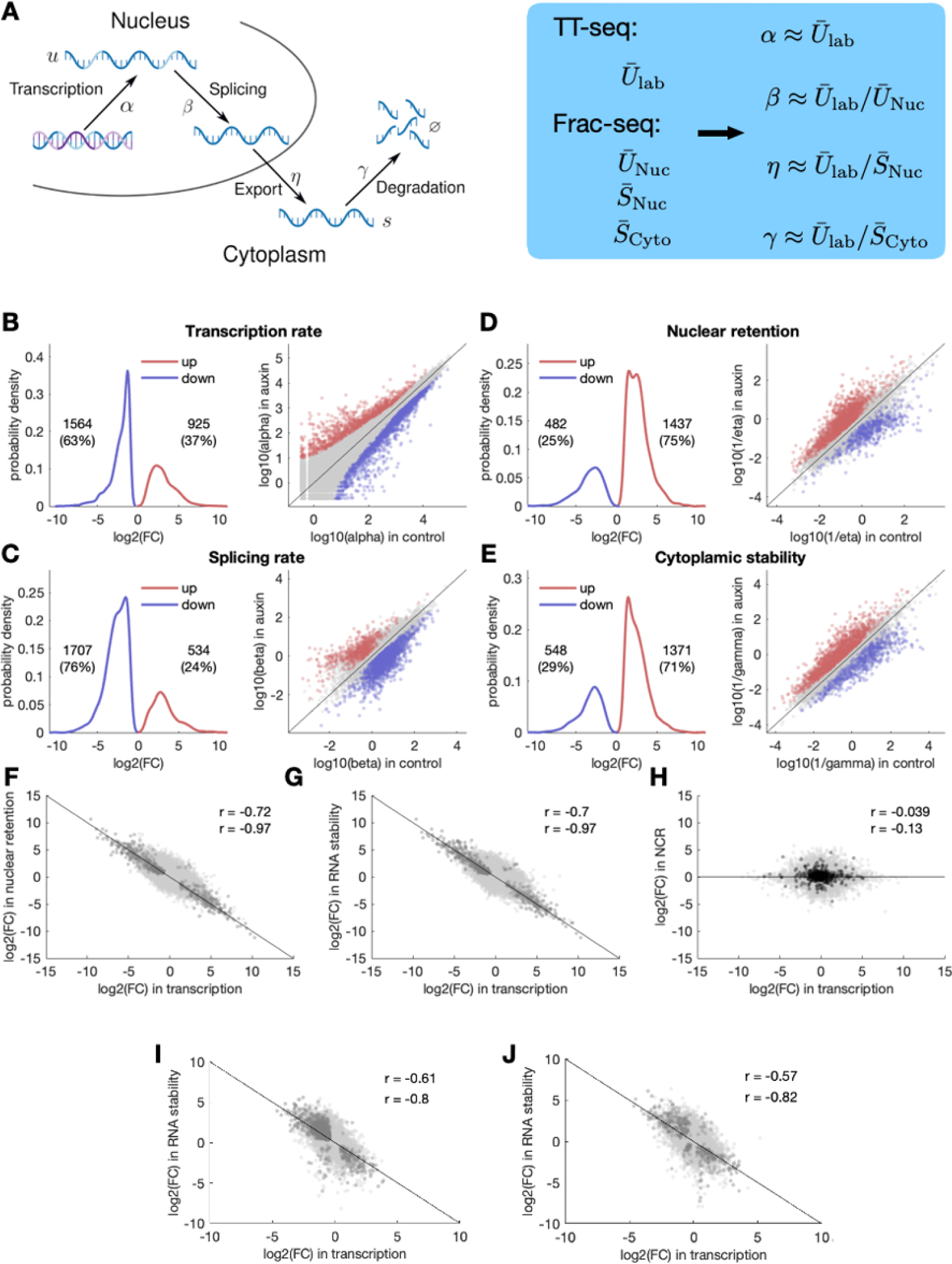
Integration of TT-seq and Frac-seq reveals gene-specific transcriptional buffering. **A:** Schematic diagram of mRNA metabolism: in the nucleus, DNA is transcribed with transcription rate *α* to unspliced pre-mRNA denoted by *u.* Introns get removed with splicing rate *²*, giving spliced mature mRNA denoted by *s*. Spliced mRNA is exported from the nucleus into the cytoplasm with an export rate *η* and is degraded in the cytoplasm with a degradation rate *γ*. Rates can be derived from TT-seq and Frac-seq data (blue box). *Ū*_*lab*_ denotes unspliced labelled RNA; *Ū_N_*, *S̄_N_* and *S̄_C_* correspond to unspliced and spliced nuclear RNA, and to spliced cytoplasmic RNA, respectively. See *Methods* for details. **B-E:** *Left*: distribution of log2 fold changes (FC) in transcription rate (*±*) (**B**), splicing rate (**C**), nuclear retention (**D**), and cytoplasmic stability (**E**) upon TIP60 depletion. Only genes with a log2 FC bigger than 1 (red) or smaller than -1 (blue) and p < 0.01 were considered. The numbers of genes that pass these criteria are shown. *Right*: scatter plot of gene-specific rates for transcription (**B**), splicing (**C**), nuclear retention (**D**), and cytoplasmic stability (**E**) in control vs. auxin. Blue and red dots represent significant up- and down-regulated genes (p < 0.01, log2 FC > |1|). Gene numbers and relative percentages in each category are indicated. **F-G:** Correlations between changes in transcription rate and changes in either nuclear retention (**F**) or cytoplasmic stability (**G**) upon TIP60 depletion. Dark dots represent genes with a significant FC (p < 0.01). Pearson correlation coefficients are displayed for all the genes (r = -0.72 and r = -0.7), and for significant genes (r = -0.97 and r = -0.97). **H:** log2 FC in transcription vs. log2 FC in nuclear to cytoplasmic ratio (NCR). Black dots represent intronless genes. Pearson correlation coefficients are displayed for all the genes (r = -0.039) or intronless genes (r = -0.13). **I-J:** Correlation between the log2 FC in transcription rate and the log2 FC in RNA stability after depletion of the ATAC subunit Zzz3 (**I**) or Yeats2 (**J**). Dark dots represent genes with a significant FC (p < 0.05). Pearson correlation coefficients are displayed for all the genes (r = -0.61 and r = -0.57) and for significant genes (r = -0.8 and r = -0.82).

Our approach allowed us to determine the rates of RNA synthesis (⍺), splicing (β), nuclear export (η), and cytoplasmic stability (γ) of *Tip60^AID^*cells treated with auxin relative to DMSO. Note that we assumed that nuclear export is typically a faster process compared to nuclear degradation (Smalec et al., 2022), and therefore nuclear retention is mainly dominated by the time mRNAs need to be translocated to the cytoplasm.

Our analyses show that upon TIP60 depletion, a greater number of RNAs exhibit reduced synthesis and splicing rates, while fewer RNAs display increased rates (**Figures 3B** and **C**). These trends are in line with the role of TIP60 in promoting transcription and underscore the tight coupling between transcription and splicing processes (Herzel et al., 2017; Ding and Elowitz, 2019). Conversely, we observed an opposing pattern for nuclear retention and cytoplasmic stability: a larger proportion of RNAs increase their nuclear and cytoplasmic half-lives in TIP60-depleted cells compared to control cells (**Figure 3D** and **E**). Strikingly, at the level of individual RNAs, the fold changes in transcription were inversely proportional to the fold changes in nuclear retention and cytoplasmic degradation (Figure **3F** and **G**). This led to the maintenance of N/C ratios for most RNAs despite alterations in their synthesis rates (Figure **3H**).

Gene-specific compensatory changes in RNA stability in response to altered RNA synthesis may be a specific response to TIP60 depletion. To test this, we compared changes in RNA abundance (measured with RNA-seq) to changes in RNA synthesis (measured with TT-seq) after inactivation of the Ada-two-A-containing (ATAC) complex, an unrelated transcriptional co-activator essential for ESC survival. The ATAC core subunits YEATS2 and ZZZ3 fused to AID were depleted by addition of auxin for 24 h (Fischer et al., 2021). We found that under these conditions, changes in RNA synthesis due to ATAC perturbation also led to compensatory gene-specific changes in RNA stability (Figure **3I-J** and Supplementary Figure **S8**). Together, these observations indicate a strong coupling between the transcription of single RNAs, their nuclear export/degradation, and their cytoplasmic decay, which we term *gene-specific buffering*.

The enhanced nuclear retention exhibited by RNAs with reduced synthesis rates in TIP60-depleted cells (shown in Figure **3F**) prompted us to investigate how transcription inhibition affects mRNA nuclear distribution in ESCs. To do so, we imaged polyadenylated RNA by fluorescence in situ hybridisation (FISH) after a 2-hour treatment with RNA polymerase II inhibitors. This treatment resulted in significant shifts in the subnuclear distribution of nuclear mRNA, which concentrated in nuclear speckles (Supplementary Figure **S9**). Thus, a reduction of mRNA synthesis prompts the rapid redistribution of mRNA to nuclear speckles in ESCs.

### Buffering is less efficient for genes not bound to TIP60

Perfect buffering is expected to maintain constant RNA levels after perturbation of transcription. However, TIP60 depletion results in alterations in the levels of some RNAs (Figure 2), suggesting that those genes are inefficiently buffered. To examine buffering efficiency, we overlaid RNA abundance data (derived from RNA-seq) onto plots correlating changes in RNA synthesis against changes in nuclear retention and cytoplasmic stability. Furthermore, we separately analysed TIP60 target genes (Figure **4A-C**) and non-target genes (Figure **4D-F**). Notably, RNA-seq data were not used to generate the biophysical model, and therefore serve as an independent validation.

**Figure 4.**
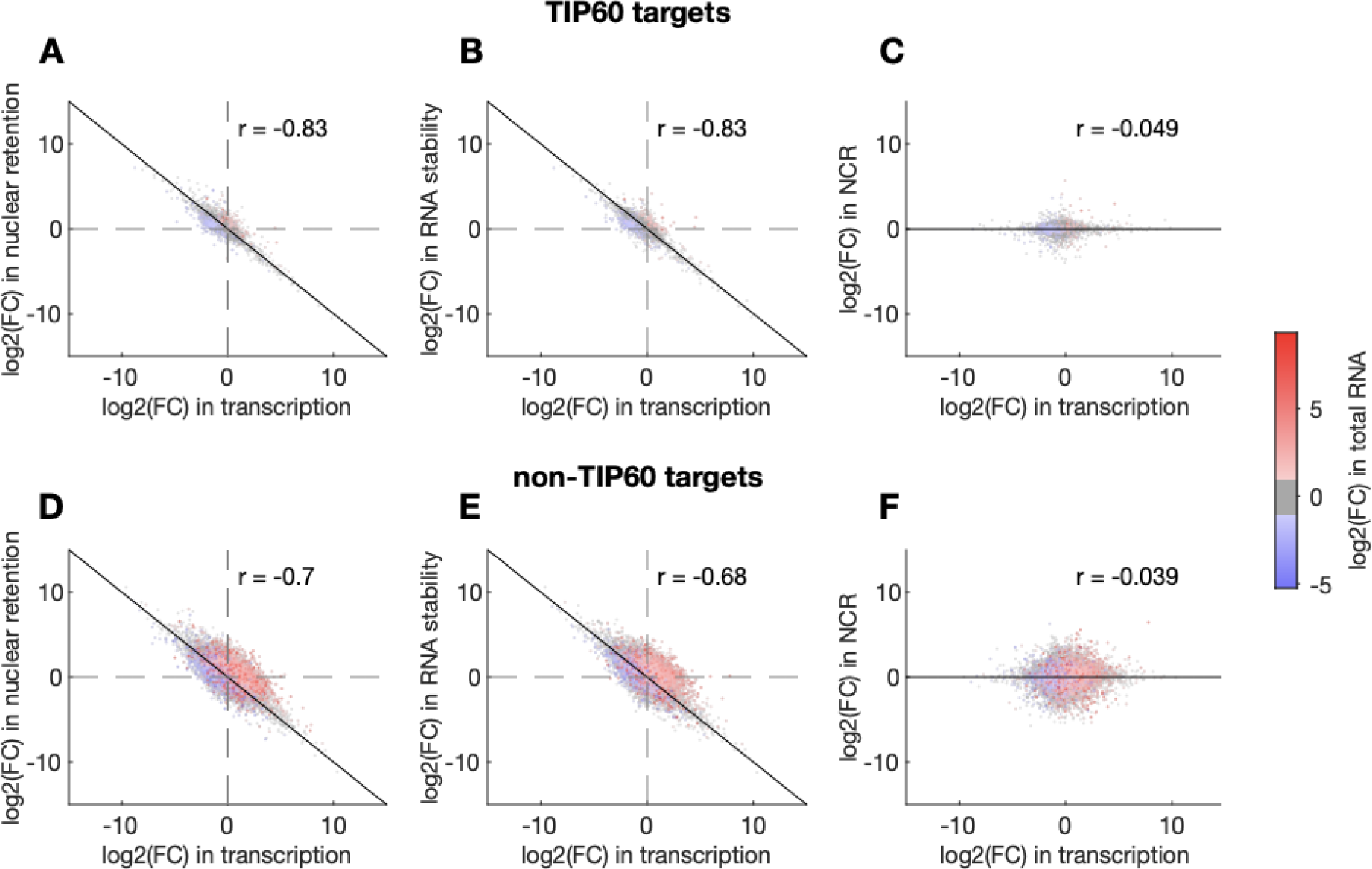
Buffering efficiency for Tip60 target and non-target genes. Correlation between changes in transcription and nuclear retention (**A** and **D**), cytoplasmic stability (**B** and **E**) and N/C ratio (**C** and **F**) after TIP60 depletion as in Figure 3F-H, for TIP60 target and non-target genes. Changes in total RNA levels are colour-coded as indicated.

This analysis reveals that most RNAs with significant changes in transcription in TIP60-depleted cells (log2 FC>|1|, p<0.01) are efficiently buffered, in both TIP60 target and non-target genes (r=0.99; grey dots in Figure **4A-B** and **4D-E**). In contrast, most changes in RNA abundance are associated with relatively small but simultaneous increases (or decreases) in both transcription and nuclear retention / cytoplasmic stability. RNAs exhibiting these changes are identifiable as, respectively, red or blue dots located away from the diagonal in Figure **4A-B** and **4D-E**. Intriguingly, changes in RNA abundance are more common in non-TIP60 transcriptional targets. Thus, the majority of gene expression changes after TIP60 depletion, which occur in genes not directly bound by TIP60, are associated with inefficient coordination between RNA synthesis and RNA export / stability.

## Discussion

The TIP60 KAT was initially identified as a transcriptional coactivator, able to acetylate various transcription factors and histone proteins (Sterner and Berger, 2000; Squatrito et al., 2006; Sapountzi et al., 2006). However, RNA sequencing of ES cells in which *Tip60* was downregulated by siRNA suggested that TIP60 mostly acts as a transcriptional repressor (Fazzio et al., 2008; Chen et al., 2013). In contrast, our results show that TIP60 actively promotes the transcription of a substantially larger set of genes compared to those it represses. Yet, this coactivator role is masked by compensatory post-transcriptional adjustments on RNA abundance.

These findings lead to two major conclusions. Firstly, the major role of TIP60 in RNA synthesis in ESCs is to act as a transcriptional coactivator for its specific target genes. This is supported by the enrichment of TIP60 targets among genes exhibiting reduced transcription in TIP60-depleted cells, while genes displaying increased transcription did not display a direct interaction with TIP60. Consequently, the role of TIP60 in repressing transcription may be of an indirect nature. For instance, TIP60 may modulate the activity and/or expression of other transcriptional regulators or RNA binding proteins governing these genes. Supporting this possibility, TIP60 associates with transcriptional repressors including lysine deacetylases (Chen et al., 2013; Xiao et al., 2003).

Secondly, the discrepancy observed between alterations in RNA abundance and synthesis rates reveals a gene-specific homeostasis mechanism that counterbalances changes in the synthesis of specific RNAs by adjusting their abundance in both the nucleus and cytoplasm. In particular, RNAs exhibiting reduced synthesis rates show increased retention in the nucleus and increased stability in the cytoplasm, while those with increased transcription display reduced residence in these compartments. Retention of mRNAs in the nucleus is associated with their recruitment to nuclear speckles, which may shield these mRNAs from degradation. Consistent with this hypothesis, we and others (Tokunaga et al., 2006) observe redistribution of mRNA to nuclear speckles after inhibition of transcription. Notably, accumulation of mRNAs in nuclear speckles has also been observed after depletion of RNA export factors, including the nuclear pore component TPR (Shibata et al., 2002; Aksenova et al., 2020).

Our observations are reminiscent of the known interdependence between global mRNA synthesis and degradation in yeast and animal cells, where inhibition of one of these processes leads to upregulation of the other, a phenomenon termed “transcript buffering” (Timmers and Tora, 2018; Hartenian and Glaunsinger, 2019). Transcript buffering is thought of as a global sensing mechanism responsive to changes in overall RNA synthesis. However, we observe simultaneous and opposite effects on RNA nuclear residence and cytoplasmic stabilisation: an increase in the stability of RNAs with reduced synthesis rates and a decrease in the stability of RNAs with increased synthesis rates. Our results suggest that buffering is ensured at the level of individual genes and may not only respond to overall transcription rates. This conclusion has significant implications for our understanding of buffering mechanisms. Specifically, our data cannot be explained by models postulating that the activity of the transcriptional machinery regulates general RNA export and/or degradation activities (Timmers and Tora, 2018; Hartenian and Glaunsinger, 2019).

The nature of the gene-specific buffering signal remains unclear. Our observation that transcriptional changes caused by perturbing two different coactivator complexes (TIP60 and ATAC) are efficiently buffered indicates that these coactivators themselves are likely not involved in the buffering mechanism. Intriguingly, promoter swap experiments in yeast have indicated that mRNA stability may be encoded within the promoter sequence, hinting at a connection between transcription and gene-specific buffering (Dori-Bachash et al., 2012). Yet, the relevance of this phenomenon to gene-specific buffering in mammalian cells has not been directly investigated. We speculate that buffering mechanisms involve the direct coupling of gene-specific RNA synthesis rates with their nuclear export and degradation rates. This coupling might be mediated by RNA “marks” that adjust RNA export and stability proportionally to RNA synthesis rates, possibly through association with RNA-binding proteins or specific RNA modifications. An example of a mark deposited proportionally to RNA synthesis rate is N6-methyladenosine (m6A), which is slightly higher in slow-transcribing mRNAs. However, this modification drives mRNA instability (Slobodin et al., 2020; Gallego et al., 2022; Lee et al., 2020b), arguing against its involvement in gene-specific buffering. The impact of other synthesis rate-dependent RNA marks on its nuclear export and stability, and their potential involvement in buffering mechanisms remains to be investigated. Additionally, exploring how buffering mechanisms are modulated during critical cellular transitions, such as embryonic stem cell differentiation, presents a fascinating area for future research.

## Acknowledgements

We thank Yves Barral, Sebastián Chávez, Pablo Navarro, Stephane Vincent, Mihaela Zavolan and members of the Mendoza, Molina and Tora laboratories for helpful discussions; Eugene Makeyev for critical reading of the manuscript, the GenomEast platform (“France Génomique” consortium ANR-10-INBS-0009); the IGBMC Imaging Centre, member of the national infrastructure France-BioImaging supported by the French National Research Agency (ANR-10-INBS-04); the IGBMC Cell Culture and Flow Cytometry Facilities, and N. Jung for cloning of CRISPR-Cas9 plasmids. This work was supported by Agence Nationale de la Recherche (ANR) CE11-0013-01_ACT, Fondation pour la Recherche Médicale (EQU-2021-03012631), and NIH MIRA (R35GM139564) grants (to LT); ANR-22-CE12-0021, Fondation ARC PGA2022010004436_4876, Eucor “The European Campus”, and the University of Strasbourg Institute of Advanced Studies USIAS-2019-046 (to MM); and by EU MSCA ITN ‘PEP-NET’ Grant Agreement n. 813282 and ANR-20-CE12-0014 (to NM). This work, as part of the Interdisciplinary Thematic Institute IMCBio+ 2021-2028 program of the University of Strasbourg, CNRS and Inserm, was also supported by IdEx Unistra (ANR-10-IDEX-0002), SFRI-STRAT’US project (ANR 20-SFRI-0012) and EUR IMCBio (ANR-17-EURE-0023) under the framework of the France 2030 Program.

## Author Contributions (CRediT author statement)

**FF**: Conceptualization, Data Curation, Investigation, Methodology, Validation, Visualization, Writing – Original Draft. **DP**: Data Curation, Formal Analysis, Methodology, Visualization. **AF**: Investigation, Methodology, Validation. **DFM**: Formal Analysis, Visualization. **KAO**: Formal Analysis. **BRSM**: Methodology, Resources. **LT**: Methodology, Resources, Funding Acquisition, Supervision, Writing – Review & Editing. **NM**: Conceptualization, Formal Analysis, Funding Acquisition, Investigation, Methodology, Visualization, Writing – Original Draft, Writing – Review & Editing. **MM**: Conceptualization, Data Curation, Funding Acquisition, Supervision, Visualization, Writing – Original Draft, Writing – Review & Editing.

## Methods

### Cell culture

Embryonic Stem Cells were grown on 0.1% gelatinized (Sigma-Aldrich, Cat# G1890) tissue-culture plates in DMEM (4.5 g/L glucose) with GlutaMAX-I supplemented with ES-tested 15% foetal calf serum (FCS, ThermoFisher Scientific, Cat# 10270-106), 0.1% β-mercaptoethanol (ThermoFisher Scientific, Cat# 31350-010), 100 U/ml penicillin, 100 μg/ml streptomycin (ThermoFisher Scientific, Cat# 15140-122), 0.1 mM non-essential amino acids (ThermoFisher Scientific, Cat# 11140-035), and 1500 U/ml leukaemia inhibitory factor (produced in-house). 3 μM CHIR99021 (Axon Medchem, Cat# 1386) and 1 μM PD0325901 (Axon Medchem, Cat# 1408) (2i) were added freshly to the medium, which was replaced every 24 h. Cells were kept in 400 µg/ml G418 for Tir1 selection. Auxin (IAA, 3-Indoleacetic acid, Sigma-Aldrich I2886) was used at 1 mM. ESC numbers were assessed using a Countess II Automated Cell Counter (Invitrogen). *Drosophila melanogaster* Schneider S2 cells (CRL-1993, ATCC) were grown in Schneider’s *Drosophila* medium (ThermoFisher Scientific, Cat# 21720-024) containing 10% FCS (heat inactivated) (Sigma-Aldrich, Cat# F7524) and 0.5% penicillin and streptomycin at 27°C.

### Plasmid construction

The plasmid expressing two gRNAs (guide RNAs, sequence: last exon, 5′-ACTGGAGCAAGAGAGGAAAG-3′; 3’ UTR, 5′-CACGAGAGCTGGCCGAACCA-3′) target the 3′ end of the endogenous the *Tip60* (*Kat5*) locus and co-express the high-fidelity Cas9 nuclease (Cas9-HF) (Kleinstiver et al., 2016). The homologous recombination (HR) template contains homology arms of approximately 800 bp surrounding the AID-FLAG-BioTag-P2A-EGFP construct. This allows TIP60 detection via the Flag tag and BirA-dependent biotinylation of the Biotin acceptor peptide, and identification of positive clones via fluorescence of free EGFP owing to the P2A self-cleaving peptide. All gRNA/Cas9-HF and HR plasmids were generated through Golden Gate cloning (Engler et al., 2009).

### Generation of Tip60 Auxin-Inducible Degron cell line

Tir1-BirA mouse ES cells (PGKpr:Tir1-HA-IRES-3xHA-BirA-SV40pr:NeoR) were produced as previously described (Fischer et al., 2021), and tested negative for mycoplasma contamination. CRISPR–Cas9 was used to generate mouse ES cells with TIP60 endogenously tagged with AID. Tir1-BirA mouse ES cells at a confluency of 70-80% were transfected with the plasmid constructs using Lipofectamine 2000 (ThermoFisher Scientific, Cat#11668019) following the manufacturer’s instructions. The donor plasmid (Tip60-AID-Flag-BioTag-P2A-EGFP) was linearized using a restriction enzyme before transfection and transfected together with a Cas9-containing transient plasmid in the Tir1-BirA expressing cell line. Cells were sorted by fluorescence-activated cell sorting by GFP expression two to three days after transfection. Three to five 96-well plates were seeded with one fluorescent cell per well using the BD FACSAria^TM^ II (BD Biosciences), following the manufacturer’s instructions. The primer pairs that were used for genotyping were the following: forward, 5′- GAGCCCCCTGTCCTTTCCTATTATG--3′; reverse, 5′- AAGGGAGATGGTAGGTTTGGGGTGAGGGCAGTAGC-3′.

### Cell cycle analysis

Cell cycle profiles were determined by labelling of propidium iodide (PI) and followed by flow cytometry analysis. Cells were fixed with 70% ethanol, 30% PBS at -20°C for at least 30 min. After being washed once with PBS, cells were treated with RNase A (Sigma-Aldrich Cat# R6513, 50 µg/ml in PBS) for 30-60 min. Cells were incubated with PI (Sigma-Aldrich Cat# P4864, 25 µg/ml in PBS) at 37°C for 20 min. The samples were analysed by flow cytometry using a MACSQuant flow cytometer (Miltenyi Biotec Inc.); data were visualised and quantified using FlowJo (Becton Dickinson).

### Whole-cell protein extraction

Cells were harvested and washed twice with 1x PBS. The cell pellet was resuspended in 1 volume of whole-cell extract buffer (50 mM Tris-HCl pH 7.9, 25% glycerol, 0.2 mM EDTA, 0.5 mM DTT, 5 mM MgCl_2_, 600 mM KCl, 0.5% NP40 and 1x protein inhibitor cocktail) and incubated for 10 min at 4°C. The salt concentration was neutralised by adding 3 volumes of IP0 buffer (25 mM Tris-HCl pH 7.9, 5% glycerol, 5 mM MgCl_2_, 0.1% NP40, 1 mM DTT and 1x protein inhibitor cocktail) and incubated for 10 min at 4°C. After centrifugation at 12,000 *g* for 10 min at 4°C, supernatants containing proteins were collected and stored at –80°C. Protein concentrations were determined using the Bradford method, using a SmartSpec^TM^ 3000 spectrophotometer (Bio-Rad).

### Western blot analysis

Proteins boiled in Laemmli buffer were separated on NuPAGE™ 4-12% gradient Bis-Tris SDS-PAGE gels, transferred onto a Hybond-C nitrocellulose membrane (GE Healthcare), then blocked with TBST containing 5% (w/v) non-fat milk (TBSTM), for 30 mins. Membranes were incubated overnight in primary antibodies diluted 1:1000 in TBS-Tween containing 1% (w/v) non-fat milk (or 1% BSA for Streptavidin-HRP), at 4°C. Membranes were washed three times with TBST, incubated in HRP-conjugated anti IgG secondary antibodies (Cell Signaling Technology) in TBSTM at room temperature, followed by further three washes with TBST. The membranes were developed using the Pierce^TM^ ECL Western Blotting Substrate (SuperSignal™ West Pico PLUS Chemiluminescent Substrate) and visualised using a ChemiDoc Imaging System (Bio-Rad). Streptavidin Protein fused to HRP (Thermo Fisher, 21126) was used to detect the TIP60-AID-Flag-BioTag fusion protein. Actin was detected with anti–β-actin (A5316; Sigma-Aldrich) antibodies.

### Total RNA extraction

Total RNA extraction was performed using TRI Reagent® (Molecular Research Center Inc., Cat# TR 188), following the manufacturer’s instructions. DNase I treatment was performed to prevent genomic DNA contamination using the TURBO DNA-free™ Kit (ThermoFisher Scientific, Cat# AM1907) manufacturer’s instructions.

### 4sU metabolic labelling

5×10^7^ cells from three independent cultures were treated with Auxin or DMSO. 24 h after Auxin treatment, the nucleoside analogue 4-Thiouridine (Glentham Life Sciences, Cat# GN6085) was added to a final concentration of 500 mM for a 10-min pulse at 37°C and 5% CO_2_. After labelling, cells were washed with ice-cold 1x PBS and immediately lysed using TRI Reagent® (Molecular Research Center Inc., Cat# TR 188).

### Purification of newly synthesised RNA

Newly synthesised RNAs were purified as previously described in detail (Rädle et al., 2013; Schwalb et al., 2016; Rabani et al., 2011). Briefly, 4sU-labelled total RNA of spike-in cells (*D. melanogaster*) was added to 250 μg of labelled total RNA from mouse ESCs in a ratio 1:10 prior to newly synthesised RNA purification. The RNA was precipitated and resuspended in 130 μl RNase-free water (Sigma-Aldrich, Cat# 95284) and sonicated on a E220 Focused-ultrasonicator (Covaris) using the following settings: 1% duty factor, 100 W, 200 cycles per burst, 80 s, to obtain fragment size range from 10 kb to 200 bp. For purification, the fragmented total RNA was incubated for 10 min at 60°C and immediately chilled on ice for 2 min to open secondary RNA structures. Biotinylation was performed in labelling buffer (10 mM HEPES–KOH pH 7.5 and 1 mM EDTA) and 0.2 mg/mL Biotin-HPDP (ThermoFisher Scientific, Cat# 21341) for 3 h at room temperature at 24°C in the dark and with gentle agitation. Unbound Biotin-HPDP was removed by adding an equal volume of chloroform/isoamyl alcohol (24:1) at 16000 *g* for 5 min at 4°C. RNA was precipitated at 20,000 *g* for 20 min with a 1:10 volume of 5 M NaCl and an equal volume of 100% isopropanol. The pellet was washed with an equal volume of 75% ethanol and precipitated again at 20,000 *g* for 10 min. The pellet was resuspended in 100 μL RNase-free water. Biotinylation and purification of 4sU-labelled RNAs was performed as described (Dölken et al., 2008; Wachutka et al., 2019). Biotinylated RNA was captured using 100 μl of streptavidin-coated μMACS magnetic beads (Miltenyi Biotec, Cat# 130-074-101) for 90 min at 24°C under gentle agitation. The μMACS columns (Miltenyi Biotec, Cat# 130-074-101) were placed on a MACS MultiStand (Miltenyi Biotec) and equilibrated with washing buffer (100 mM Tris–HCl pH 7.5, 10 mM EDTA, 1 M NaCl, 0.1% Tween 20) twice on the columns before adding the samples. The columns were then washed once with 600 μl, 700 μl, 800 μl, 900 μl and 1 ml with washing buffer. Flow-through was collected for recovery of unlabeled preexisting RNA. RNA-4sU was eluted with two washes of 100 μL of freshly prepared 100 mM dithiothreitol (DTT). Reverse Transcription (RT) was performed with 1 μg total RNA and using 0.2 μg random hexamer primers (ThermoFisher Scientific, Cat# SO142) and 200 U SuperScript® IV Reverse Transcriptase (ThermoFisher Scientific, Cat# 18090050) following manufacturer’s instructions.

### qPCR

Real-time quantitative PCR reactions were performed using a LightCycler 480 system (Roche) with SYBR Green 2× PCR Master Mix I (Roche, Cat# 04887352001) and 1 μM of forward and reverse primer respectively. The primer pairs used for qPCR are listed in Supplementary Table **T1**. Relative gene expression was calculated based on the obtained threshold values using the 2^−(ΔΔCT)^ method (Pfaffl, 2001).

### RNA Frac-Seq

Nuclear and cytoplasmic RNAs were purified as previously described in detail (Lee et al., 2020a). Optimising cell fractionation techniques for ESCs enabled the isolation of nuclear and cytoplasmic fractions from control and TIP60-depleted cells, followed by the purification and sequencing of their respective RNA content. Briefly, cells growing in three 150 mm plates at 60-80% confluency were harvested with trypsin followed by centrifugation at 12000 rpm for 5 min, then the supernatant was discarded and the cell pellet washed 3 times with 1x PBS. Subsequently, 10% of cell volume was subjected as the total RNA fraction and the remaining (90%) was resuspended in 500 μL φ buffer [150 mM potassium acetate, 5 mM magnesium acetate, 20 mM HEPES pH 7.4, 1 mM sodium fluoride, 1 mM sodium orthovanadate, 25× protease inhibitor cocktail (Roche), 1:1000 dilution of SUPERase In™ RNase (Invitrogen) and 0.1% diethylpyrocarbonate]. Then, 500 μl of φ buffer containing 1% Triton X-100 (Thermo Scientific) and 0.2% sodium deoxycholate was gently added to the resuspended cells and incubated for 3 min on ice. After that, the cell sample was centrifuged at 12000 rpm for 5 mins. Finally, cytoplasmic RNAs were extracted from the supernatant, and the nuclear RNAs were extracted from the pellet using TRI Reagent® (Molecular Research Center Inc., Cat# TR 188) manufacturer’s instructions. The same RNA extraction procedure was performed to extract RNA from the ‘total’ fraction. DNase I treatment was performed to prevent genomic DNA contamination using the TURBO DNA-free™ Kit (ThermoFisher Scientific, Cat# AM1907), following the manufacturer’s instructions.

### Library preparation and sequencing

Total RNA-Seq libraries were generated from 500 ng of total RNA. Before cDNA synthesis, cytoplasmic and mitochondrial ribosomal RNA (rRNA) were removed using a biotin-streptavidin magnetic bead-based procedure with the riboPOOL kit targeting HMR ribosomal rRNA (siTOOLs Biotech, Planegg/Martinsried, DE) according to manufacturer’s instructions. Total RNA-seq libraries were then generated using TruSeq Stranded mRNA Library Prep kit and TruSeq RNA Single Indexes kits A and B (Illumina, San Diego, CA) omitting the poly-A selection step and starting from the fragmentation step. Total RNA-Seq libraries were generated from 50 ng of total RNA for the TT-seq experiment and from 600 ng of total RNA for the Frac-seq experiment, using Illumina Stranded Total RNA Prep, Ligation with Ribo-Zero Plus kit and IDT for Illumina RNA UD Indexes, Ligation (Illumina, San Diego, USA), according to manufacturer’s instructions. Abundant ribosomal RNAs were depleted by hybridization to specific DNA probes and enzymatic digestion. Briefly, for these three experiments, the depleted RNA was fragmented into small pieces using divalent cations at 94°C for 2 minutes. Cleaved RNA fragments were then copied into first strand cDNA using reverse transcriptase and random primers followed by second strand cDNA synthesis using DNA Polymerase I and RNase H. Strand specificity was achieved by replacing dTTP with dUTP during second strand synthesis. The double stranded cDNA fragments were blunted using T4 DNA polymerase, Klenow DNA polymerase and T4 PNK. A single ’A’ nucleotide was added to the 3′ ends of the blunt DNA fragments using a Klenow fragment (3′ to 5′exo minus) enzyme.

The cDNA fragments were ligated to double stranded adapters using T4 DNA Ligase. The ligated products were enriched by PCR amplification (30 sec at 98°C; [10 sec at 98°C, 30 sec at 60°C, 30 sec at 72°C] x 12 cycles (13 cycles for TT-seq); 5 min at 72°C). Surplus PCR primers were further removed by purification using AMPure XP beads (SPRIselect beads for TT-seq and Frac-seq) (Beckman-Coulter, Villepinte, France) and the final cDNA libraries were checked for quality and quantified using capillary electrophoresis. All the libraries were sequenced with 2 x 100 base pairs on an Illumina HiSeq 4000 sequencer or on an Illumina NextSeq 2000 sequencer for the Frac-seq experiment. Image analysis and base calling were carried out using RTA v.2.7.3 and bcl2fastq v.2.17.1.14.

### Sequence analysis total RNA-seq, TT-seq and Frac-seq

Reads were preprocessed using CUTADAPT v.1.10 (Martin, 2011) in order to remove adaptors and low-quality sequences and reads shorter than 40 bp. rRNA sequences were removed for further analysis. Reads were mapped onto the mm10 assembly of the *Mus musculus* genome using STAR v.2.5.3a (Dobin et al., 2013). For TT-seq data, due to the spike-in, reads were mapped onto a hybrid genome composed of *Mus musculus* and *Drosophila melanogaster*. Gene expression was quantified from uniquely aligned reads using HTSeq-count v.0.6.1p1(Anders et al., 2014) with annotations from Ensembl release 102 and union mode (and ‘-t gene’ for TT-seq data in order to take into account reads aligned onto introns). Only non-ambiguously assigned reads have been retained for further analyses. Comparisons of interest have been performed using R 3.3.2 with DEseq2 version 1.16.1 (Love et al., 2014). More precisely, read counts were normalised from the estimated size factors using the median-of-ratios method and a Wald test was used to estimate the P-values. P-values were then adjusted for multiple testing with the Benjamini and Hochberg method (Benjamini and Hochberg, 1995). For TT-seq data, size factors were estimated using *Drosophila* spike-in. To determine if a category of mRNA was more impacted by the depletion of Tip60, genes with a mean FPKM > 1 were considered and genes from the mitochondrial genome were excluded. For protein coding genes or lincRNA, bins were created according to the median pre-mRNA length (<10, 10–20, 20–30, 30–40, 40–50, >50 kb) or according to the number of exons (1, 2, 3, 4, 5, 6, 7, 8 for protein coding genes and 1, 2, 3, 4, 5 for lncRNA). DESeq2 log_2_ fold changes (with fold change shrinkage) from auxin-treated vs control comparison were compared between bins using Mann-Whitney-Wilcoxon tests with Benjamini-Hochberg adjustment for multiple comparisons. For Frac-seq data, the difference of DESeq2 log_2_ fold changes (log_2_(Nuclear/Cytoplasmic) _auxin-treated_ - log_2_(Nuclear/Cytoplasmic) _Control_) was used for the comparisons. Genes having a length <10 kb (or one exon) were compared to genes from other length bins (or other exon bins). The same approach was used to determine if the canonical or non-canonical histones were impacted by the depletion of Tip60.

### ChIP-seq analysis

FASTQ files were retrieved from GEO for p400IP_WT and IgG (Fazzio et al., 2015). After read mapping onto mm10 mouse genome using Bowtie2 v.2.4.5, BAM files were converted into BED files using BEDTools v.2.30.0. Blacklisted regions, available on this link: https://github.com/Boyle-Lab/Blacklist/tree/master/lists, were removed from the BED files. Peak calling was performed using MACS2 v.2.7.1 and IgG was used as control. Peak annotation was carried out using annotatePeaks.pl program from HOMER v.4.11, with Ensembl version 102 as annotation file.

### Mathematical modelling of RNA metabolism

In order to integrate TT-seq and Frac-seq data, we propose a biophysical model to describe mRNA accumulation in both the nucleus and the cytoplasm, extending the previous RNA velocity approach (Gaidatzis et al., 2015; La Manno et al., 2018). In particular, we assume that unspliced nuclear mRNA (*u_N_*) is synthesised with a constant transcription rate α and spliced out with a constant splicing rate β. In turn, spliced nuclear mRNA (*s_N_*) is translocated into the cytoplasm with a constant export rate η. Finally, spliced cytoplasmic mRNA (*s*_*c*_) is degraded with a constant degradation rate γ. Consequently, mRNA accumulation dynamics in the different compartments is governed by the following system of ordinary differential equations:

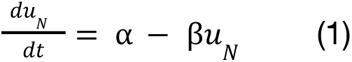

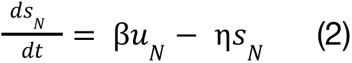

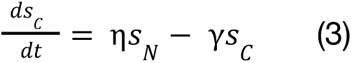

Note that the effective rates α, β, η and γ are considered gene-specific and characterise the effective speed at which complex multi-steps processes of mRNA metabolism occur. Furthermore, we assume that mRNA export is typically faster than nuclear degradation (Smalec et al., 2022), and therefore, the nuclear retention of spliced mRNA is mainly governed by mRNA translocation.

The typical mRNA half-life has been reported to be in the order of 9 hours (Schwanhäusser et al., 2011). Therefore, we assumed that cells reached a new quasi-equilibrium state after the 24-hour treatment with auxin. Consequently, we solved the model for the steady state by setting the right-hand side of the equations above to zero, yielding the following result:

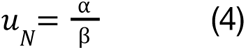

[Mth5]

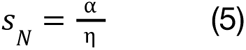

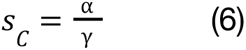

Importantly, by mapping intronic and exonic reads from the Frac-seq experiments, we were able to obtain estimates for the levels of nuclear unspliced and spliced mRNA, as well as cytoplasmic spliced mRNA averaged over the replica, denoted as *Ū*_N_, *S*^̄^_N_ and *S*^̄^_C_. In addition, from TT-seq experiments, we estimated the labelled unspliced mRNA from TT-seq averaged over replica, *Ū*_*lab*_. Plugging these quantities into equations (4), (5) and (6) and assuming that labelled RNA is a good readout of transcription rate, we obtain the following gene-specific estimates for the key rates of RNA metabolism up to a scaling factor:

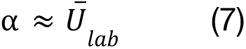

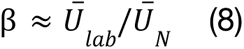

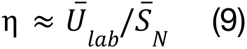

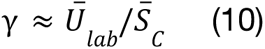

Finally, we calculated errors for each rate estimate based on the standard errors of unspliced and spliced mRNA levels by applying error propagation.

For depletion of the ATAC subunits Zzz3 and Yeats2, TT-seq data were obtained from (Fischer et al., 2021) and RNA-seq was performed under the same conditions. In the absence of Frac-seq data, RNA degradation rates were estimated from equation (10) in which total spliced RNA levels in whole cells were used instead of cytoplasmic levels. Gene-specific rates are provided in Supplementary Table **T2** (WT and TIP60 depletion) and Supplementary Table **T3** (WT, YEATS2, and ZZZ3 depletion).

### Brightfield microscopy

Images were acquired on a ZEISS Axio Observer.Z1/7 Brightfield microscope with a Plan-Apochromat 63× objective using ZEN 3.3 (blue edition) acquisition software.

### Immunofluorescence with poly(A) RNA FISH

Coverslips were coated at 37°C with 0.1% gelatin (Sigma-Aldrich, Cat# G1890) for 1 h. Cells were plated on the pre-coated coverslips and grown for 24 h, treated with 1mM Auxin or DMSO, followed by fixation using 4% paraformaldehyde (PFA) (Electron Microscopy Sciences, 15710) in PBS for 15 min, followed by 3 PBS washes. Permeabilization of cells was performed using 0.5% Triton X-100 (Sigma-Aldrich, X100) in PBS for 10 min, followed by 3 PBS washes. Cells were blocked with 5% BSA (MP Biomedicals, Cat# 160069) for 1 h at room temperature. Cells were then incubated with the primary antibody (anti SC-35/SRRM2, ab11826, Abcam) at a concentration of 1:200 overnight in PBS at 4°C. Cells were washed with PBS 3 times, followed by incubation with secondary antibodies at a concentration of 1:500 in PBS for 1 h. After washing twice with PBS, cells were fixed using 4% PFA in PBS for 10 min. Poly(A) RNA FISH was performed as described (Tsanov et al., 2016). Briefly, after two washes of PBS, cells were incubated with hybridization buffer (15% formamide from Sigma-Aldrich in 1× SSC) for 15 min, and then overnight at 37°C in the hybridization buffer containing 1 μM of the poly(A) probe for 100 µl of the final volume, 0.34 mg/ml tRNA, 2 mM VRC (Sigma-Aldrich), 0.2 mg/ml RNase-free bovine serum albumin (BSA) (Molecular Biology Grade), and 10% dextran sulphate (Sigma-Aldrich, Cat# D8906). The next day, samples were washed twice for 30 min in the hybridization buffer at 37°C. The nuclei were stained in 1 mg/ml DAPI for 60 min. Cells were then washed once in PBS, followed by mounting the coverslip onto glass slides with ProLong Gold (Invitrogen, P36934). Images were acquired on a Leica confocal microscope with a 63× objective using LAS X acquisition software (Leica).

### Image analysis

Images derived from a single confocal Z-slice were subjected to semi automated post-processing using FIJI software and a custom macro. Nuclei were initially segmented by applying the Otsu method to the DAPI channel, resulting in a mask outlining nuclear contours. The mean fluorescence intensity within these contours was then quantified in the Poly(A)+RNA channel for each nucleus. Subsequently, nuclear mRNA speckles were identified by thresholding the Poly(A)+RNA channel using the MaxEntropy method. A Gaussian blur (sigma=1) was applied to the resulting mask, followed by a secondary threshold using the Otsu method to generate a mask specific to RNA particles. Particles outside the nuclear contours were excluded, and size and circularity filters were applied to eliminate artefacts. Visual assessment was employed during data processing to ensure accurate nuclear and foci segmentation, with specific measurements being discarded if the semiautomated segmentation method exhibited poor performance. For each biological replicate, the area and mean fluorescence values of nuclei and mRNA speckles were normalised by the median value of the untreated condition (DMSO), yielding fold change values. These values were then pooled together for subsequent statistical analysis. The pooled data underwent a Kruskal-Wallis test followed by Dunn’s multiple comparisons test (comparing DMSO vs. Triptolide and DMSO vs. Flavopiridol), which was calculated using Graphpad Prism.

## Data sources and availability

The RNA sequencing datasets are available on the Gene Expression Omnibus (GEO) under the GSE253313 accession number at this link: https://www.ncbi.nlm.nih.gov/geo/query/acc.cgi?acc=GSE253313

## Supplementary Figures and Tables

**Supplementary Figure S1.**
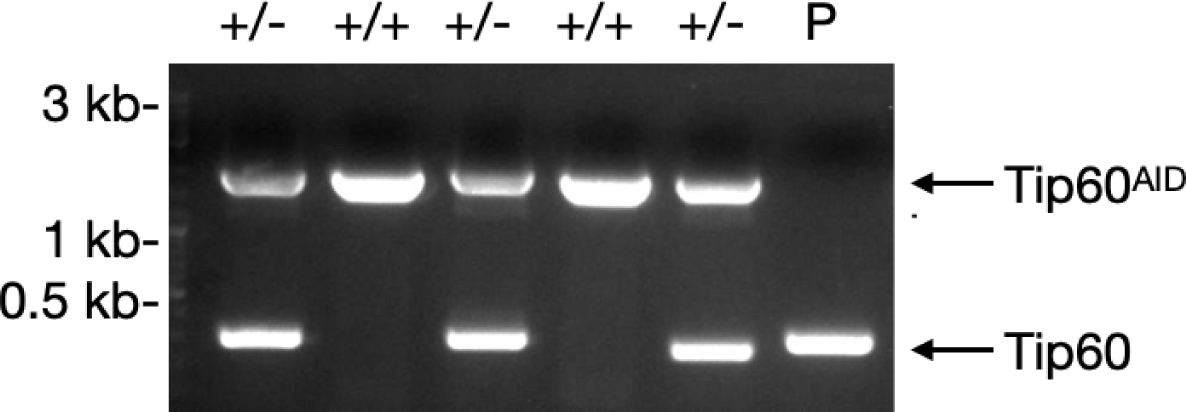
Genomic PCR of independent mESC clones demonstrating the integration of the sequence AID-FLAG-BioTagP2A-EGFP into genomic loci of *Kat5* (Tip60) gene.

**Supplementary Figure S2.**
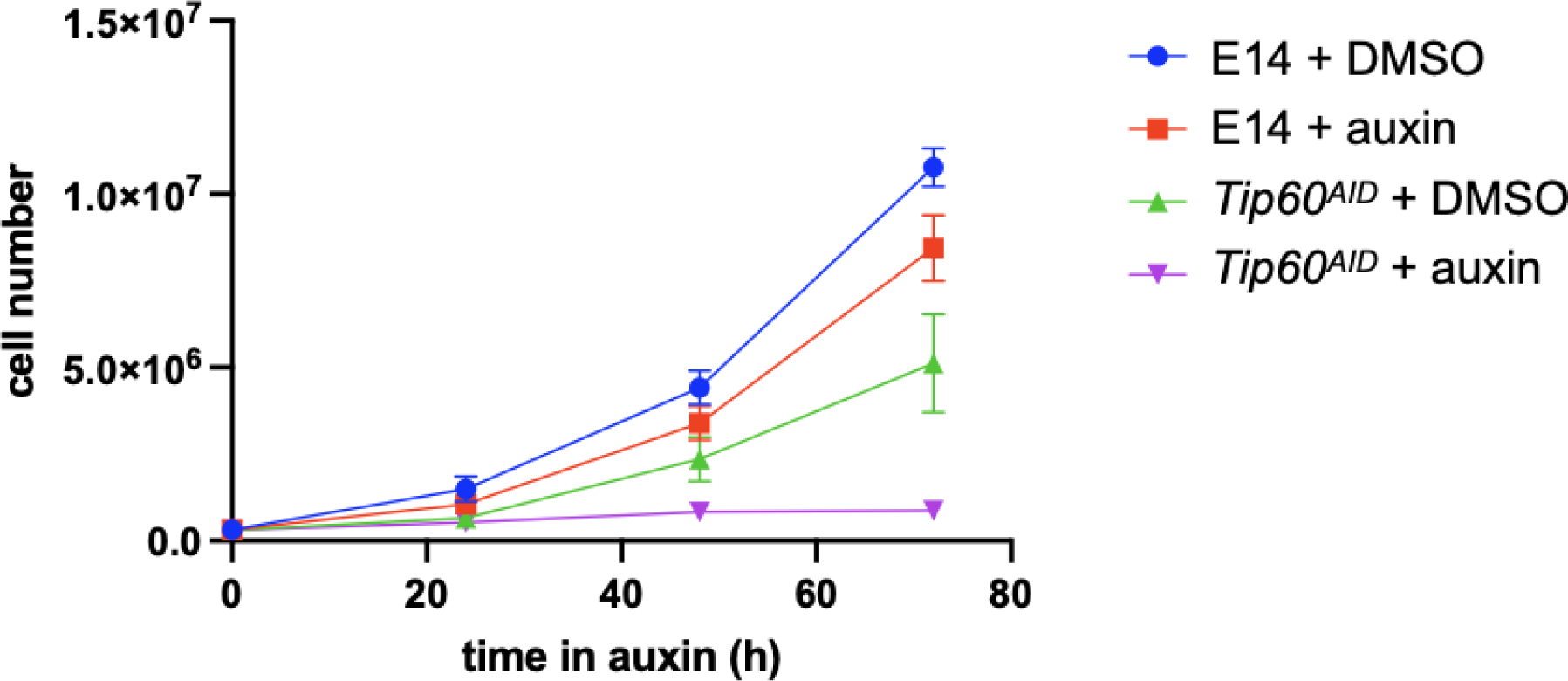
Cell number (mean and SEM of n=3 independent experiments) of the indicated cells grown in LIF medium and treated with DMSO or 1 mM auxin.

**Supplementary Figure S3.**
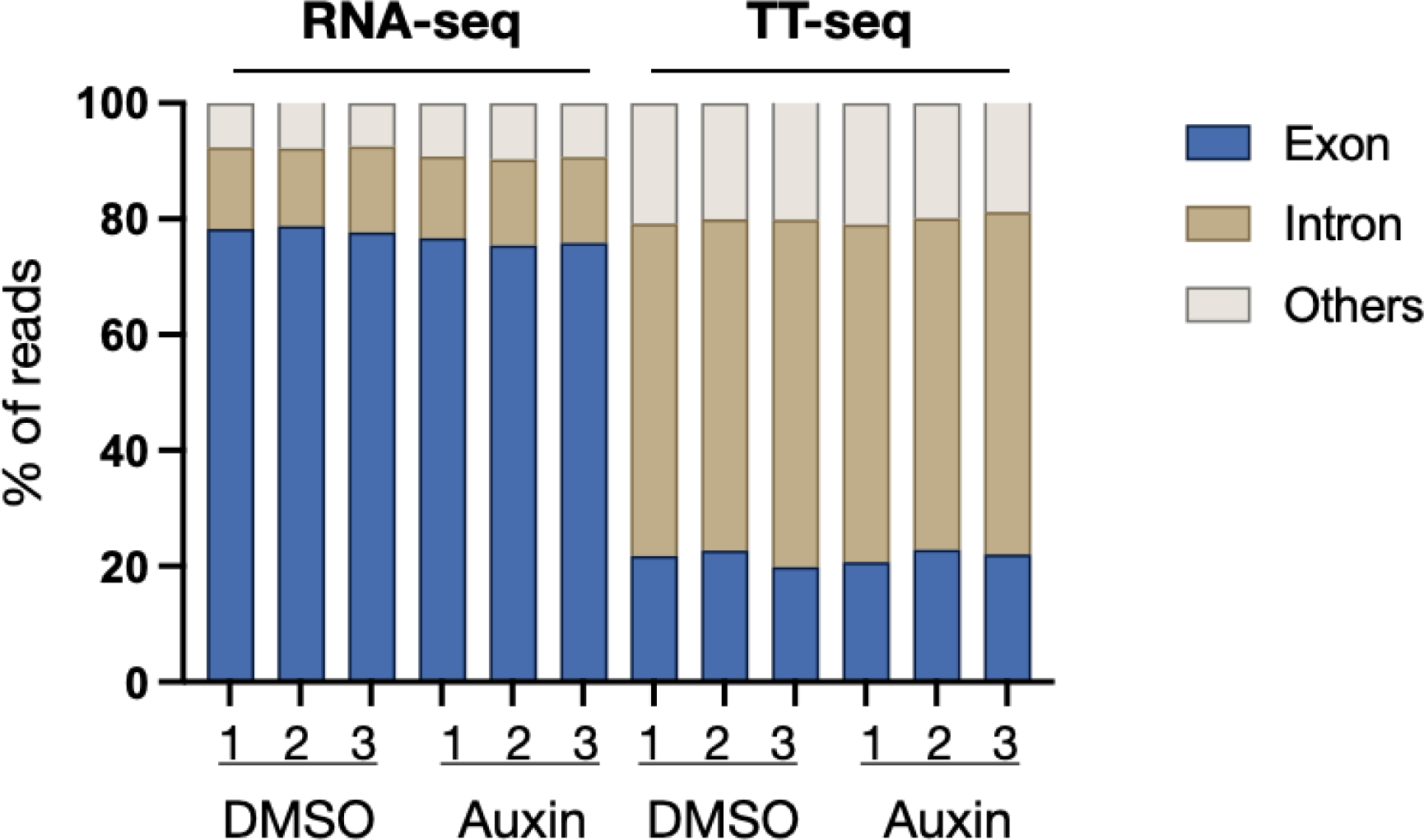
Proportion of reads mapped to the indicated genomic elements for independent replicates (1-3) of RNA-seq and TT-seq experiments. Besides reads aligning against exons and introns, reads matching exon-intron junctions, exons-intergenic junctions and intergenic regions are represented as “others”.

**Supplementary Figure S4.**
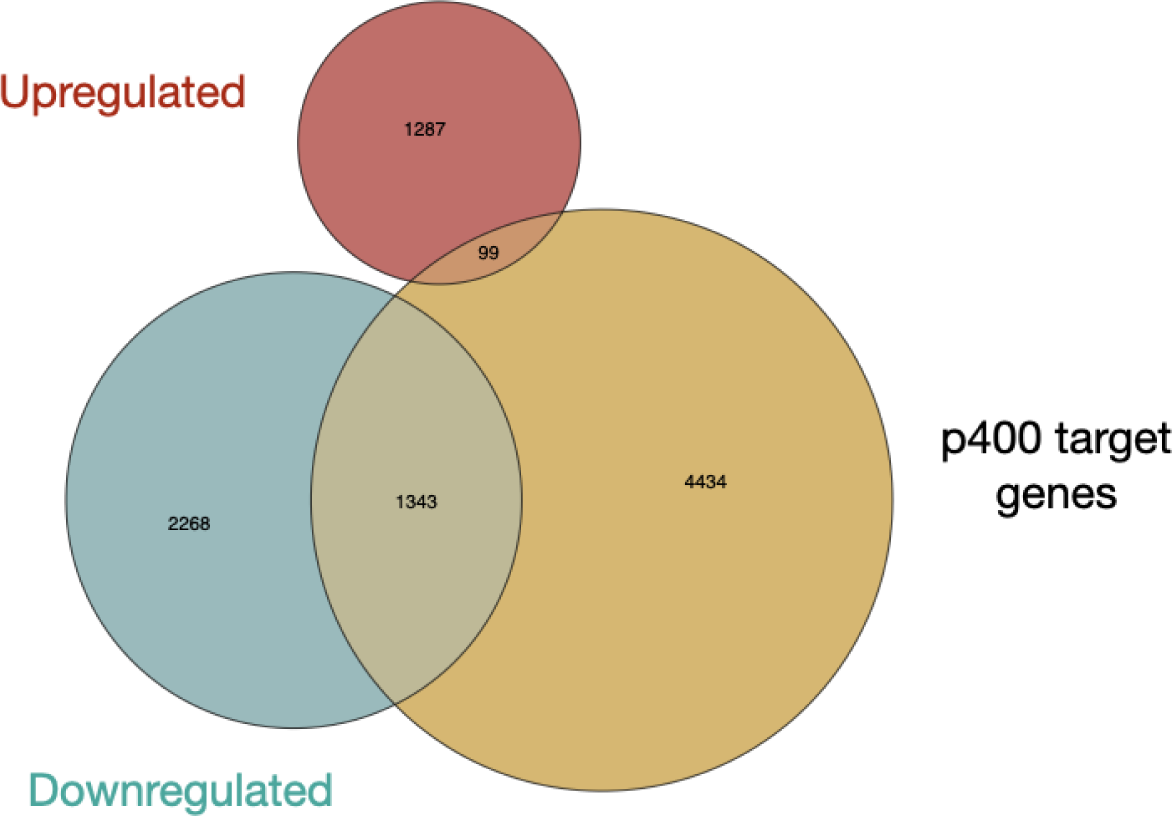
Venn diagram designating the overlap between differentially expressed genes in DMSO vs auxin-treated *Tip60^AID^* cells and p400-associated genes (Chen et al., 2015) assessed by TT-seq.

**Supplementary Figure S5.**
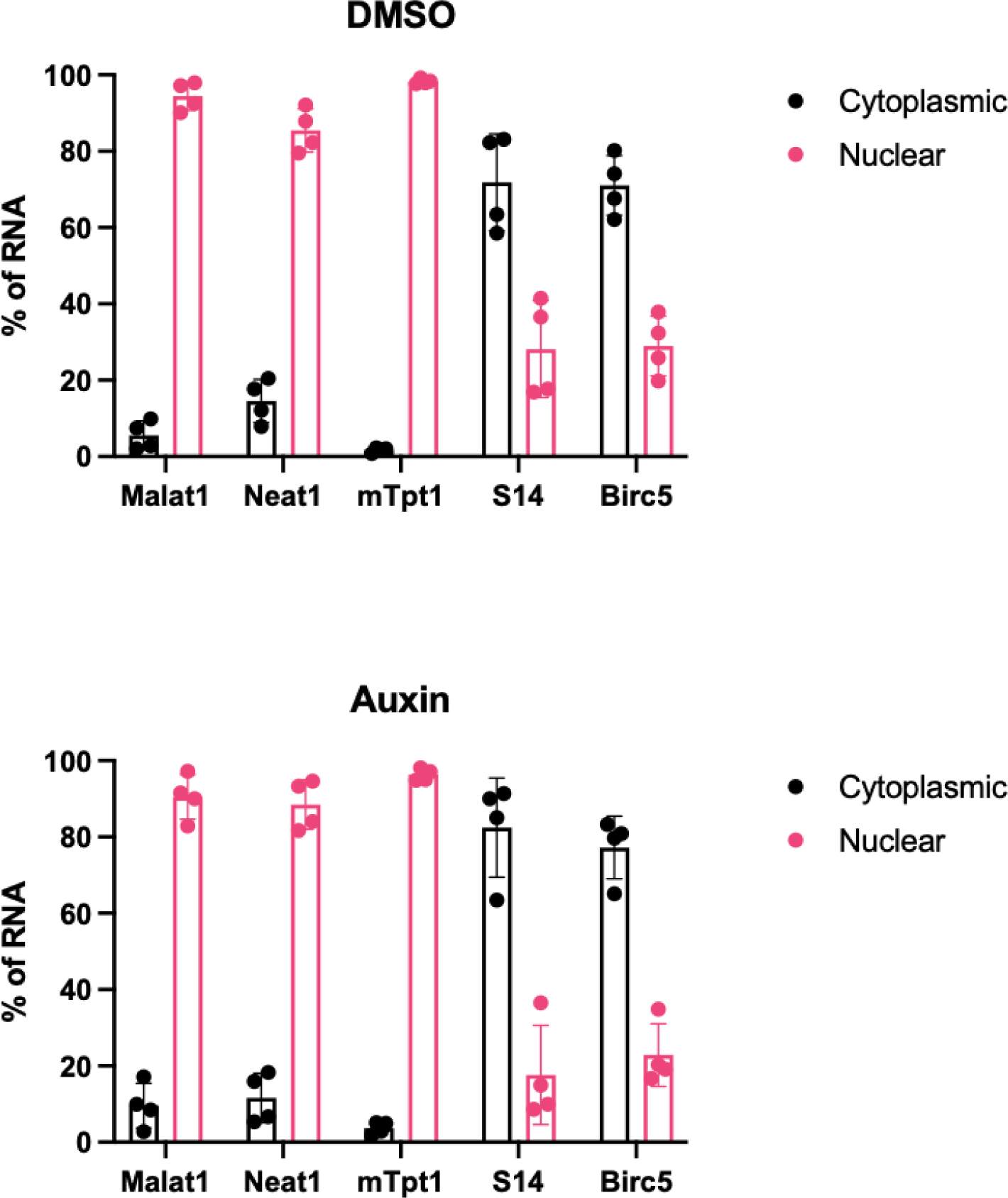
RT-qPCR analysis of the indicated RNAs from nuclear and cytoplasmic fractions (mean and SD of n=4 biological replicates). Malat1 and Neat1 are nuclear lncRNAs. mTpt1 corresponds to an intronic region. Both RPS14 (S14) and BIRC5 mRNAs are expected to localise within the cytoplasm.

**Supplementary Figure S6:**
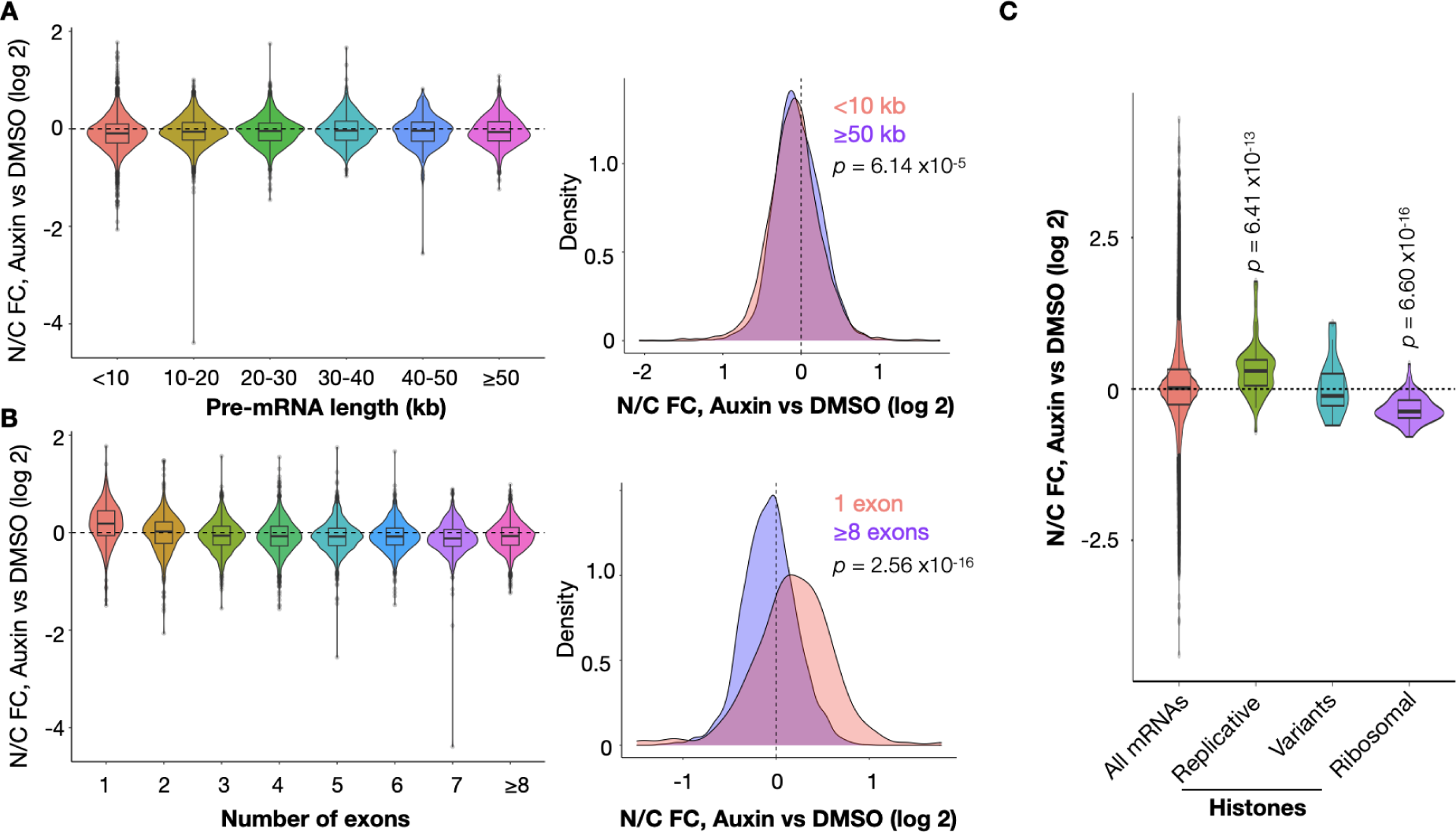
TIP60 depletion affects the N/C ratio of specific RNA categories. **A**: Violin plot of TIP60-dependent changes in nucleo/cytoplasmic ratio (auxin v DMSO fold change) for pre-mRNAs of varying lengths in 10 kb windows, and density plot showing that very short (<10 kb) and very long (>50 kb) pre-mRNAs are not significantly enriched in the nucleus after Tip60 depletion. **B**: Violin and density plots as in (A), but for pre-mRNAs containing various numbers of introns. Note that intronless pre-mRNAs show a slightly higher nuclear accumulation after TIP60 depletion. **C**: Violin plot as in (A-B) for specific mRNA classes. Note that replicative histone mRNAs are slightly enriched in the nucleus of TIP60-depleted cells, whereas ribosomal protein genes are slightly enriched in the cytoplasm.

**Supplementary Figure S7.**
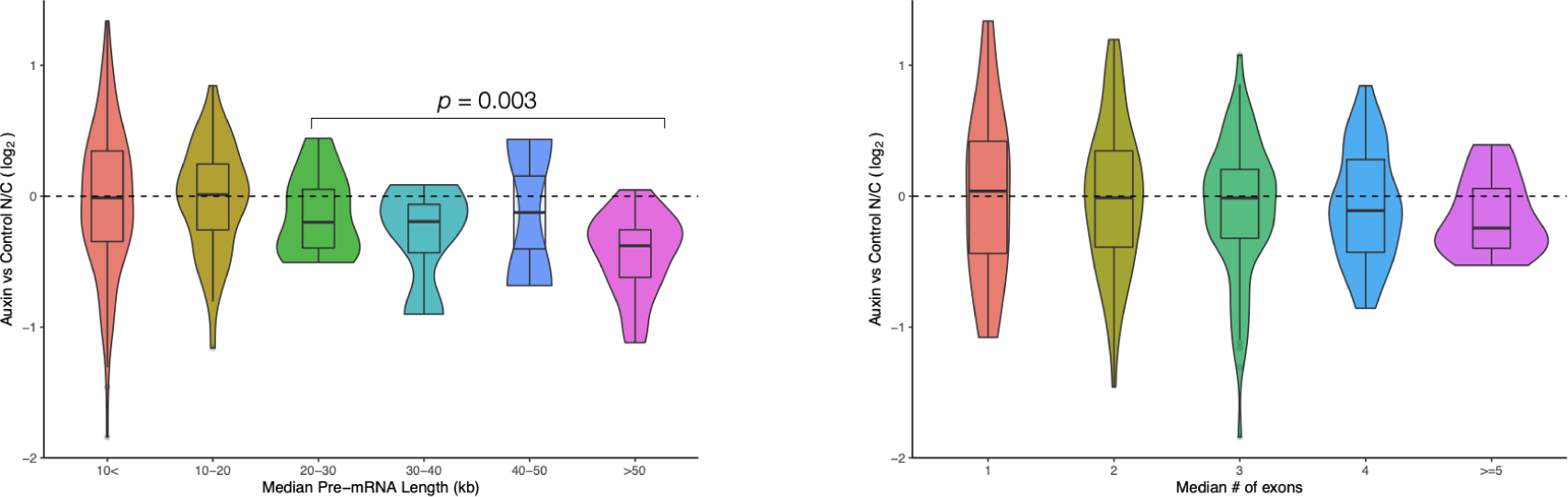
Violin plot of TIP60-dependent changes in nucleo/cytoplasmic ratio (auxin v DMSO fold change) for lncRNAs of varying lengths in 10 kb windows (left) and for lncRNAs containing various numbers of introns. Note that lncRNAs of length >50 kb show a slightly higher cytoplasmic accumulation after TIP60 depletion.

**Supplementary Figure S8:**
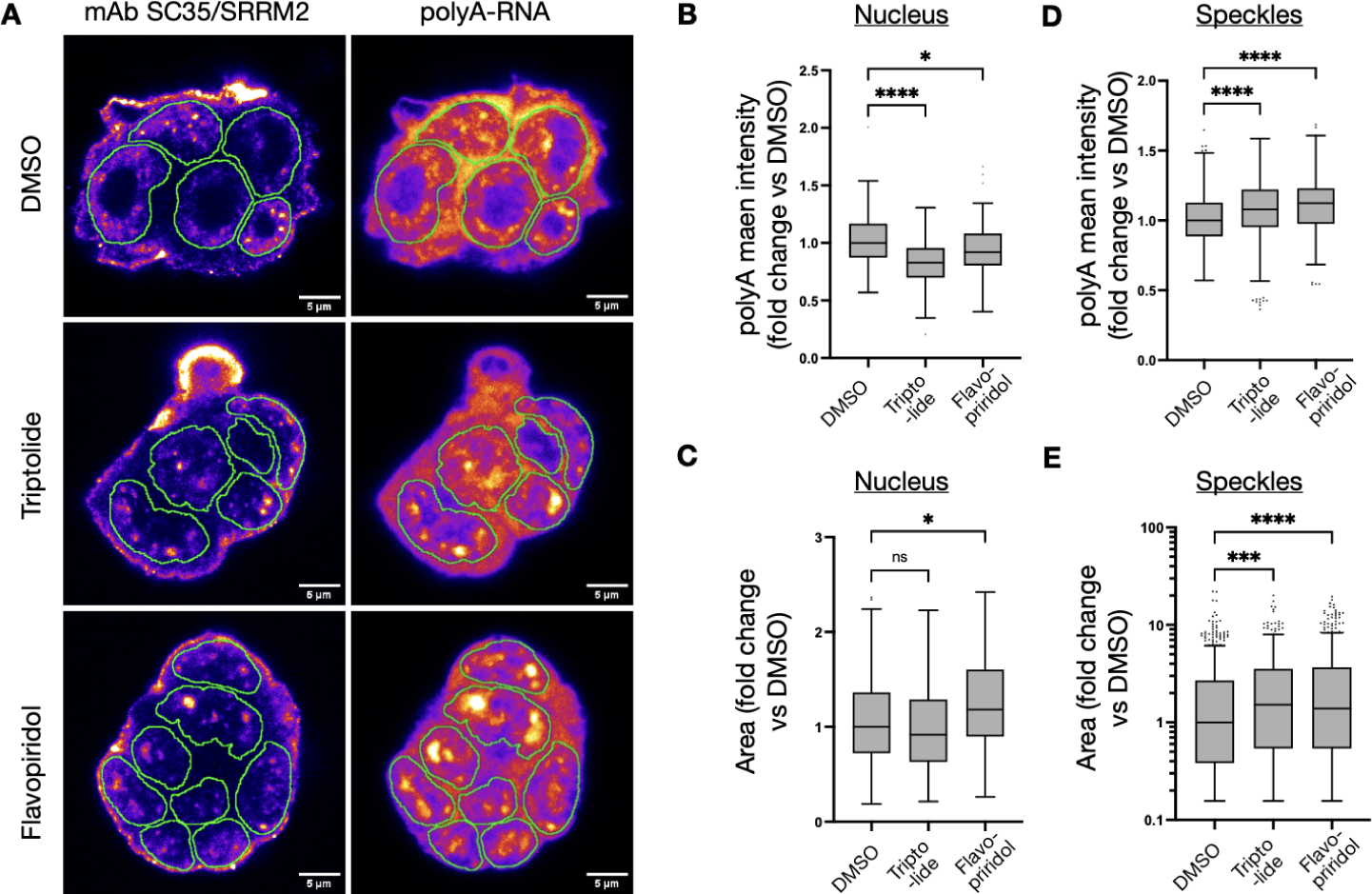
Nuclear redistribution of mRNA upon inhibition of transcription. (**A**) Representative images of mouse embryonic stem cell colonies treated with DMSO, Triptolide, or Flavopiridol for 2 hours. Cells were fixed and stained with DAPI to label nuclei, SC35/SRRM2 monoclonal antibody labelling nuclear speckles, and Cy3-tagged polyT oligonucleotide labelling mRNA. Nuclear contours are depicted in green. Scale bar, 5 μm. (**B-E**) Box plots showing the fold change compared to the DMSO condition for the following parameters: (**B**) nuclear polyA mean intensity, (**C**) nuclear area, (**D**) polyA mean intensity on speckles, and (E) speckle area (note the logarithmic scale on the y-axis). Whiskers were calculated using the Tukey method. ns, not significant (p > 0.05); * p ≤ 0.05; ** p ≤ 0.01; *** p ≤ 0.001; **** p ≤ 0.0001. Cells were pooled from two independent experiments.

**Supplementary Figure S9:**
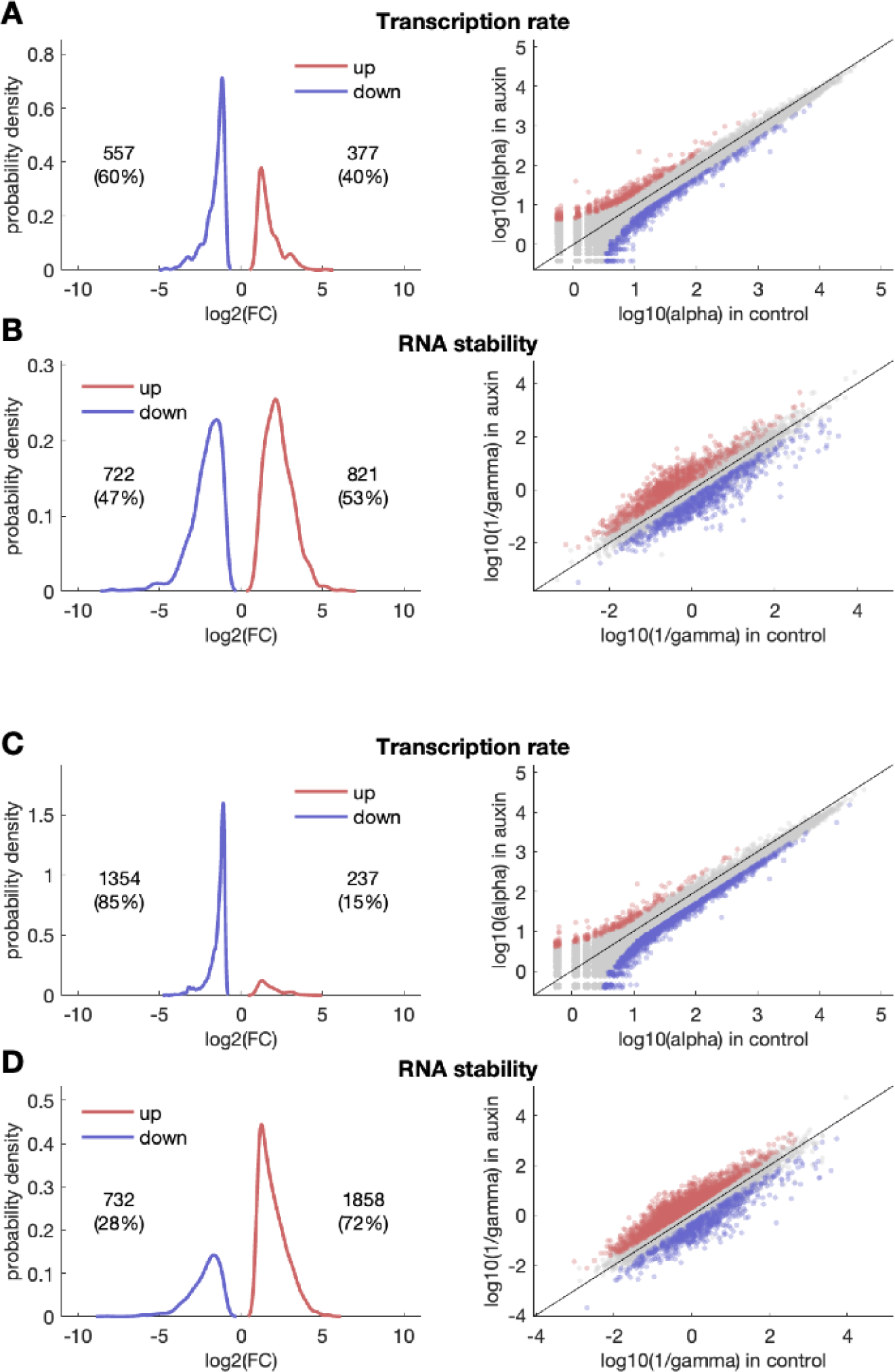
A-B: Integration of TT-seq and total RNA-seq of ZZZ3 depletion experiments. *Left*: distribution of log2 fold changes (FC) in transcription rate (*α*) (**A**) and RNA stability (**B**) upon Zzz3 depletion. Only genes with a log2 FC bigger than 1 (red) or smaller than -1 (blue) and a p-value < 0.05 were considered. The numbers of genes that pass these criteria are shown. *Right*: scatter plot of gene-specific rates for transcription (**A**) and RNA stability (**B**) in control vs. auxin. Blue and red dots represent significant (p-value < 0.01) upregulated genes (log2 FC > 1) and downregulated genes (log2 FC < -1). Gene numbers and relative percentages in each category are indicated. **C-D:** similar plots as above for the integration of TT-seq and total RNA-seq of YEATS2 depletion experiments.

**Supplementary Table T1.**
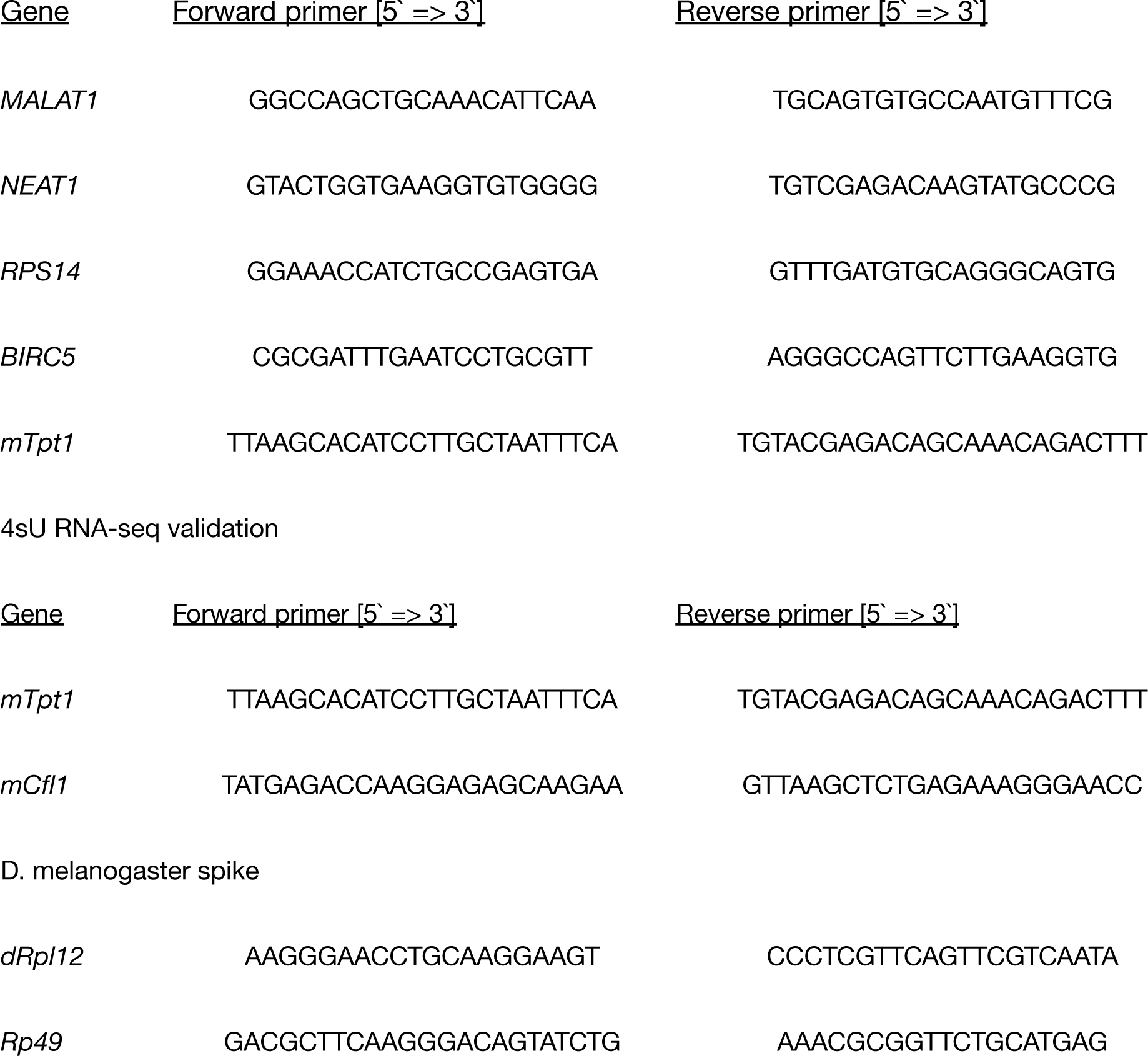
RT-qPCR primers used.

